# ORIGIN OF THREE RELATED MOVEMENT GENE MODULES IN PLANT VIRUSES: EVOLUTIONARY RADIATIONS OF BENYVIRUS-LIKE RNA REPLICONS

**DOI:** 10.1101/2022.01.19.477018

**Authors:** Sergey Y. Morozov, Andrey G. Solovyev

## Abstract

Previous studies have shown that the RNA genomes of some plant viruses encode two related genetic modules required for the virus movement over host body, containing 2 or 3 genes and named as triple gene block (TGB) and binary movement block (BMB). In this paper, we revealed a novel related movement gene module, called Tetra-Cistron Movement Block (TCMB). It was found to be encoded by virus-like transcriptome assemblies of moss *Dicranum scoparium* and Antarctic flowering plant *Colobanthus quitensis*. These TCMBs are encoded by second RNA components of the putative two-component viruses related to plant benyviruses. Like RNA-2 of benyviruses, TCMB RNA-2 contains the 5’-terminal coat protein gene. General organization of TCMB is very similar to TGB: it includes RNA helicase gene which is followed by two small overlapping cistrons encoding hydrophobic proteins with a distant sequence similarity to TGB2 and TGB3 genes. However, TCMB includes also forth 5’-terminal gene coding for protein with an obvious similarity to double-stranded RNA-binding proteins belonging to the DSRM AtDRB-like superfamily. Finally, we suggest the proposed involvement of replicative beny-like helicases in evolution of the BMB and TCMB movement genetic modules.

## INTRODUCTION

All triple gene block (TGB)-containing viruses are represented by a large variety of plant RNA viruses within the orders *Martellivirales* (family *Virgaviridae*), *Tymovirales* (familes *Alphaflexiviridae* and *Betaflexiviridae*) and *Hepelivirales* (family *Benyviridae*). They have a positive-sense, single-stranded genome consisting of one to four RNA segments (Morozov and Solovyev, 2003; Verchot-Lubicz et al., 2010; Koonin et al., 2020). The TGB-encoded movement proteins, referred to as TGB1, TGB2 and TGB3, perform directed transport of viral genomes to and through plasmodesmata (PD) into adjacent non-infected cells. TGB1 contains protein domain of RNA helicase of superfamily 1 (SF1), whereas TGB2 and TGB3 are the small membrane-associated proteins and contain highly hydrophobic amino acid segments (Verchot-Lubicz et al., 2010). Recently, binary movement block (BMB), which is related to TGB, was found in kitaviruses (family *Kitaviridae*, order *Martellivirales*). This gene module includes only two genes (BMB1 and BMB2). Although pairs of BMB1/TGB1 and BMB2/TGB2 proteins are quite similar in structural, functional and phylogenetic aspects (Morozov and Solovyev, 2015, 2020; Solovyev and Morozov, 2017; Lazareva et al., 2017; 2021), BMB gene module lacks analog of TGB3. Importantly, recent data showed that some viruses from genera *Potexvirus* and *Carlavirus* encode no TGB3 proteins, despite the fact that TGB1 and TGB2 proteins of these viruses are significantly similar to those of potexvirus-like TGBs (Morozov and Solovyev, 2015). These observations support the hypothesis that the TGB3 gene could be a less essential component (an accessory cistron) of transport gene module. Conceivably, TGB3 could be evolved as an additional cistron if BMB was the first transport module of this type appeared in virus genomes or, alternatively, could be eliminated during evolution of earliest TGB-related movement modules (Morozov and Solovyev, 2015, 2020; Solovyev and Morozov, 2017). Taking into account these considerations, identification of new TGB/BMB-like gene modules in lower land plants could shed a new light on the early steps of TGB/BMB evolution. So far, only a single TGB-containing virus-like RNA assembly has been revealed in non-seed plants, namely, bird’s-nest fern *Asplenium nidus*. This fern virus shows gene arrangements and sequence similarities indicating its close relatedness to benyviruses (Morozov and Solovyev, 2015).

In our recent paper, we draw attention to partial transcriptome assemblies in the Antarctic flowering plant *Colobanthus quitensis*, where a probable evolutionary early variant of TGB was found (Cq-TGB) (Solovyev and Morozov, 2017). Importantly, the Cq-TGB1 protein sequence is more similar to BMB1 in comparison with TGB1 proteins (Fig. 1). Moreover, the central hydrophilic region of Cq-TGB3 protein located between two transmembrane sequence segments shows clear amino acid sequence similarity to the Cq-TGB2 protein and exhibits conservation of most amino acid residues invariant in other TGB2 proteins (Solovyev and Morozov, 2017). These data suggest that Cq-TGB could arise from a BMB-like module by duplication of hydrophobic protein gene. Importantly, the Antarctica coast flora was isolated from the rest of the world for approximately 20 million years (Cantrill and Poole, 2013). This fact provided a reasonable basis for considering Cq-TGB as one of the ancient movement gene modules and further prompted us to search available sequence data to find more viral gene blocks related to TGB/BMB modules in transcriptomes of non-seed plants and extant land plants.

**Fig. 1.**
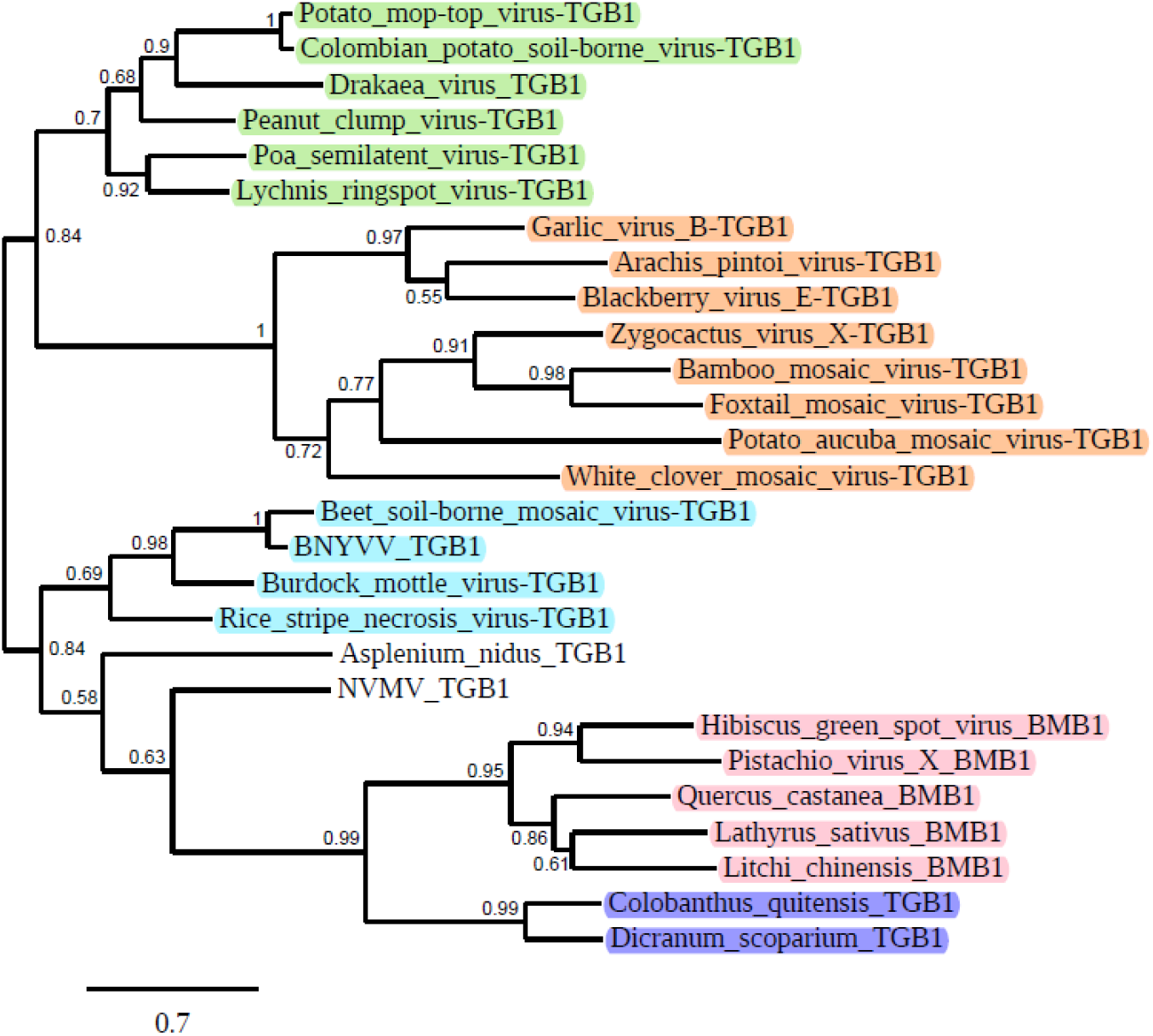
Phylogenetic analysis of the helicase domains derived from the aligned deduced amino acid sequences of the proteins encoded by TGBs and BMBs. The phylogenetic unrooted tree was constructed using the maximum likelihood method based on the amino acid sequence alignments (http://www.phylogeny.fr/simple_phylogeny.cgi). The bootstrap values obtained with 1000 replicates are indicated on the branches, and branch lengths correspond to the branch line’s genetic distances. The genetic distance is shown by the scale bar at the lower left. BNYVV – beet necrotic yellow vein virus; NVMV – Nicotiana velutina mosaic virus. Hordei-like TGB1 helicases are in green, potex-like helicases – in brown, benyvirus TGB1 helicases are in blue, BMB1 helicases found two viruses and three VLRAs (*Q. castanea, L. sativus* and *L. chinesis*) are in pink, TCMB helicases found in the corresponding VLRAs (*C. quitensis* and *D. scoparium*) are in dark blue.

In this paper, we reported new virus-like RNA assemblies (VLRAs) in the NCBI TSA and SRA databases and identified a novel plant virus movement gene module in the transcriptome samples of two wild plant species. This gene module could be classified as structurally and evolutionary related to BMB and TGB. Additionally, we presented new data supporting an idea that the movement gene modules related to TGB could initially originate in the benyvirus-like replicons.

### NOVEL MOVEMENT GENE MODULE IN THE VIRUS-RELATED PLANT TRANSCRIPTOMES: TETRA-CISTRON MOVEMENT BLOCK

We undertook a systematic analysis of RNA-seq datasets from Viridiplantae available in the NCBI open-access TSA and SRA in the end of November, 2021. Our TBLASTn search of plant TSA data collection, using Cq-TGB encoded helicase (accession GCIB01126289) as a query, resulted in the identification of a new partial VLRA (accession HANF01089872) in moss *Dicranum scoparium* (family *Dicranaceae*, order Dicranales) encoding protein with obvious similarity to Cq-TGB1 (Fig. 1). Using transcriptome sequencing data for the *D. scoparium* SRA experiment (ERX3824048), we further assembled a nearly full-length sequence of the contig (Ds-VLRA2) comprised 3,996 nt excluding the poly(A) tail. The open reading frame (ORF) prediction at ExPASy (ESTscan) showed that the contig contains six ORFs flanked by a 5’ untranslated region (5’ UTR, at least 183 nt) and a 3’ UTR (165 nt) (Fig 2A). The resulting *D. scoparium* VLRA exhibits a gene content and arrangement quite similar to that in RNA 2 of benyviruses (Fig. 2B) (Saito et al., 1996). Indeed, this RNA encodes the 5’-terminal coat protein (CP) gene (ORF1) and TGB-like module (Fig. 2A). Although the CP of Ds-VLRA2 (220 aa in length) has only marginal amino acid similarity with the members of genus *Benyvus*, it shows obvious similarity to other tobamovirus-like CPs, namely, the tobacco rattle virus CP (genus *Tobravirus*; family *Virgaviridae*) (ABE27877, 31% identity, E-value 1e-11), and the pea early-browning virus CP (genus *Tobravirus*; family *Virgaviridae*) (CAA07067, 29% identity, E-value 2e-11).

**Fig. 2.**
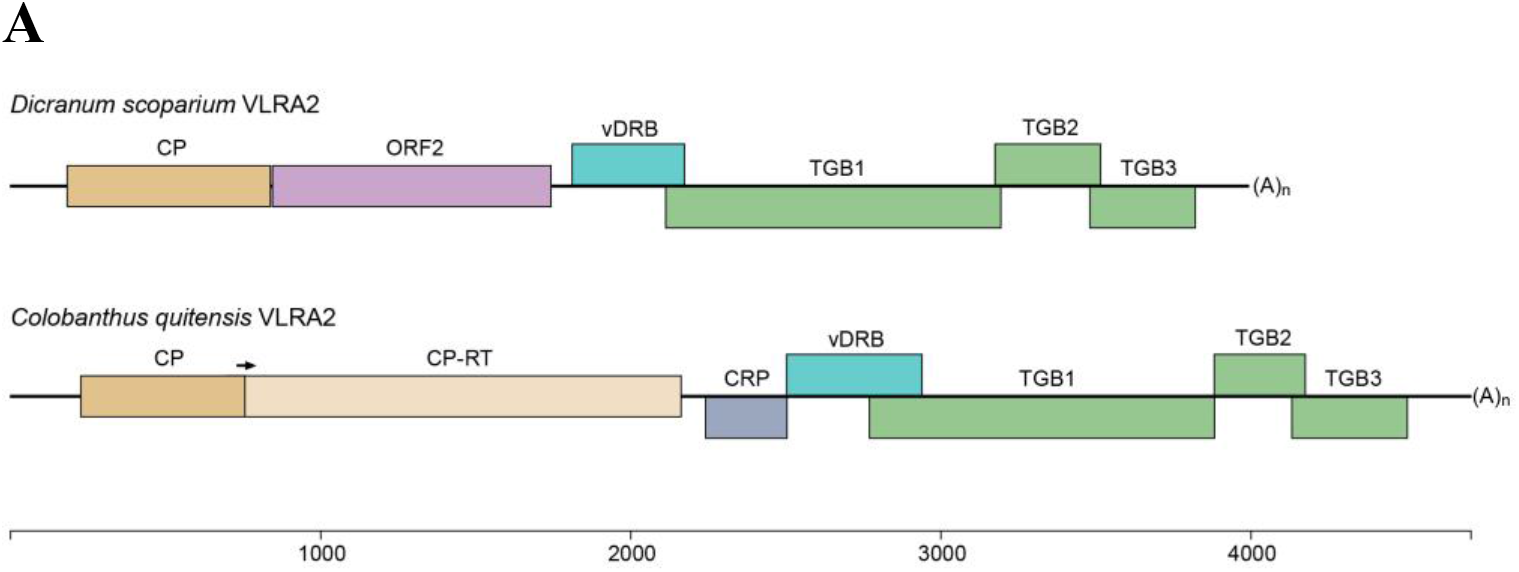
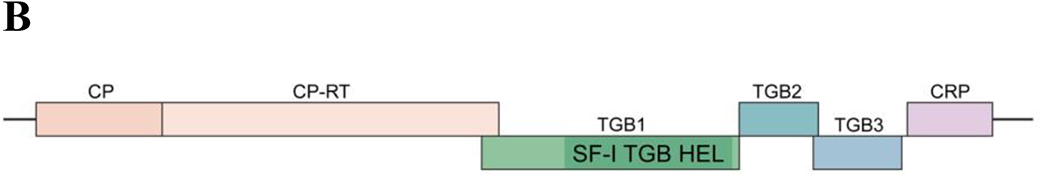
Comparison of gene organization of RNA2 genomic components encoding multicomponent cell-to-cell transport systems in *D*.*scoparium* and *C*.*quitensis* VLRAs (**A**) and Beet necrotic yellow vein virus (**B**). Genes are shown as boxes with the names of the encoded proteins. Genes of proteins potentially involved in cell-to-cell movement (TGB and vDRB) are shown in green, dark green, light green and blue-green. Genes encoding small hydrophobic proteins are shown in blue. Replicative genes are shown in yellow. Arrows indicate read-through codons in CP-RT proteins. CRP – cysteine-rich protein, CP-RT – coat protein read-through protein.

The ORF2 and ORF3 of Ds-VLRA2 encode proteins of 301 aa and 120 aa residues in length, respectively (Fig.2A). BLASTn and BLASTx analyses showed that ORF2 protein has no significant sequence homology with known virus polypeptides (data not shown). The NCBI Conserved Domain Database (CDD) analysis identified that the ORF2 protein could contain a possible domain related to the SMC superfamily (Accession No. cl34174, E-value 6.56e-03). The SMC (structural maintenance of chromosomes) domain is found in the proteins that bind DNA and act in organizing and segregating chromosomes (Lehmann, 2005). Additional protein domain analyses using ExPASy (ProtScale) software predicted a coiled-coil motif located at amino acid positions 136–185 and two highly hydrophobic sequences positioned at residues 45–63 and 279-299 in ORF2 protein (Fig. 3).

**Fig. 3.**
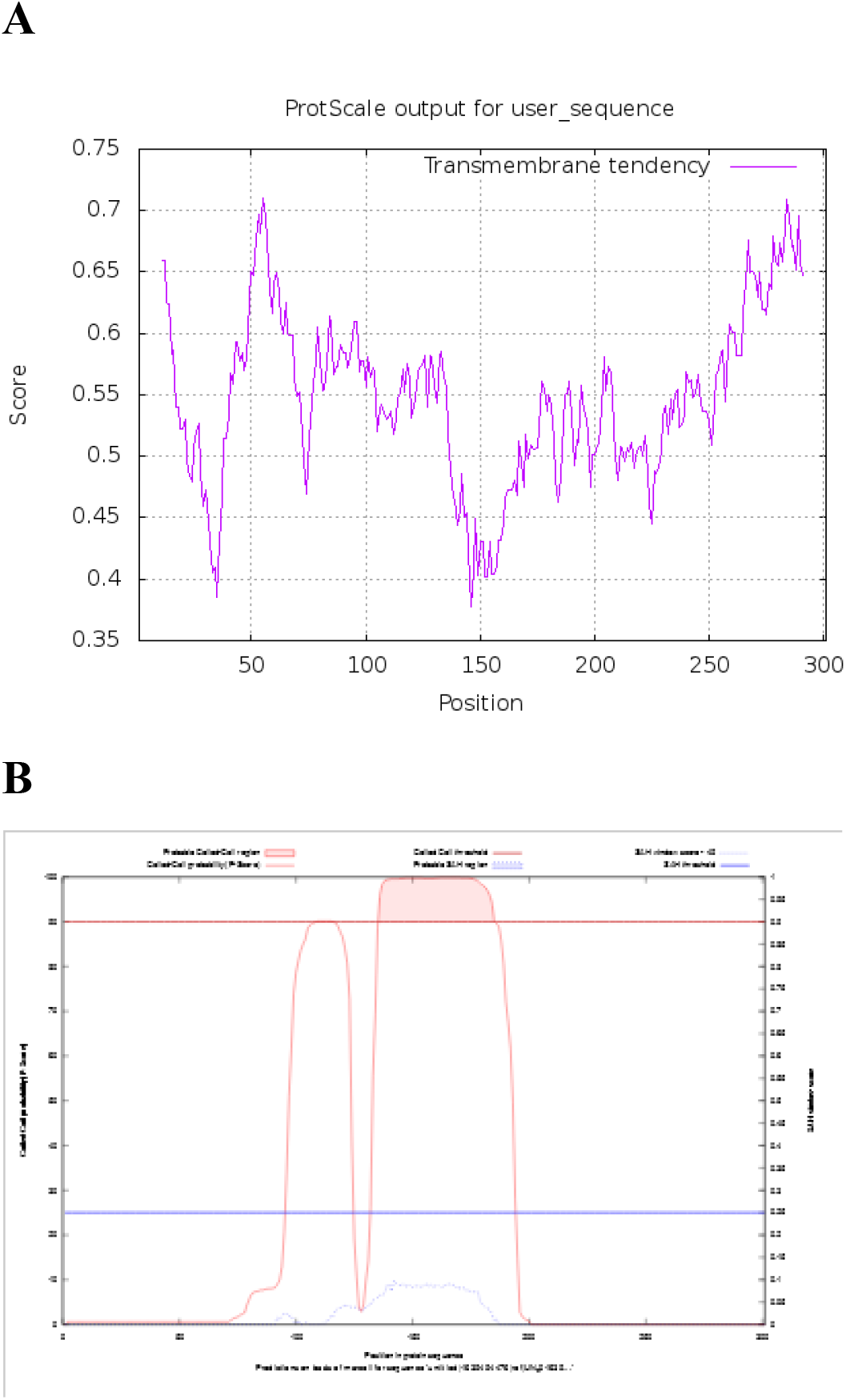
(**A**) Prediction of hydrophobic membrane-bound regions (https://web.expasy.org/protscale/) and (**B**) coiled-coil segments (above red cut-off line) (https://waggawagga.motorprotein.de/) in ORF2 protein of Ds-VLRA2.

The NCBI CDD analysis also identified a possible domain in the ORF3 protein, corresponding to the DSRM_AtDRB-like superfamily (Accession No. cl00054, E-value 1.6e-03). Therefore, the Ds-VLRA2 ORF3 protein was named viral DRB (vDRB). The DSRM protein domain superfamily is a well-known protein structural motif of 65-70 aa in length that adopts an alpha-beta-beta-beta-alpha fold and binds double-stranded RNAs (dsRNAs) of various origin and structure. This family includes a group of *Arabidopsis thaliana* double-stranded RNA-binding proteins termed AtDRB1-AtDRB5. Members of this group usually contain two DSRM domains. They bind dsRNA and are involved in RNA-mediated silencing (Han et al., 2004; Eamens et al., 2012) and/or dsRNA-triggered immunity against viruses (Fátyol et al., 2020). The vDRB protein encoded by Ds-VLRA2 contains a single DSRM showing conservation of key residues specific for DSRM_AtDRB-like proteins (Fig. 4). Interestingly, AtDRB-like proteins are encoded not only by flowering plants but also representatives of lower vascular plants (Lycopodiopsida), mosses (Bryophyta) and liverworts (Marchantiophyta). Moreover, these proteins can be revealed in present-day Charophyte algae (members of Zygnemophyceae and Charophyceae families), which are assumed to be the closest relatives of land plants descendent of the organisms that took part in initial colonization of terrestrial habitats (Fig. 4). Importantly, the Ds-VLRA2 vDRB protein, unlike AtDRB1-AtDRB5, contains a hydrophobic transmembrane segment at the N-terminus (Fig. 5).

**Fig. 4.**
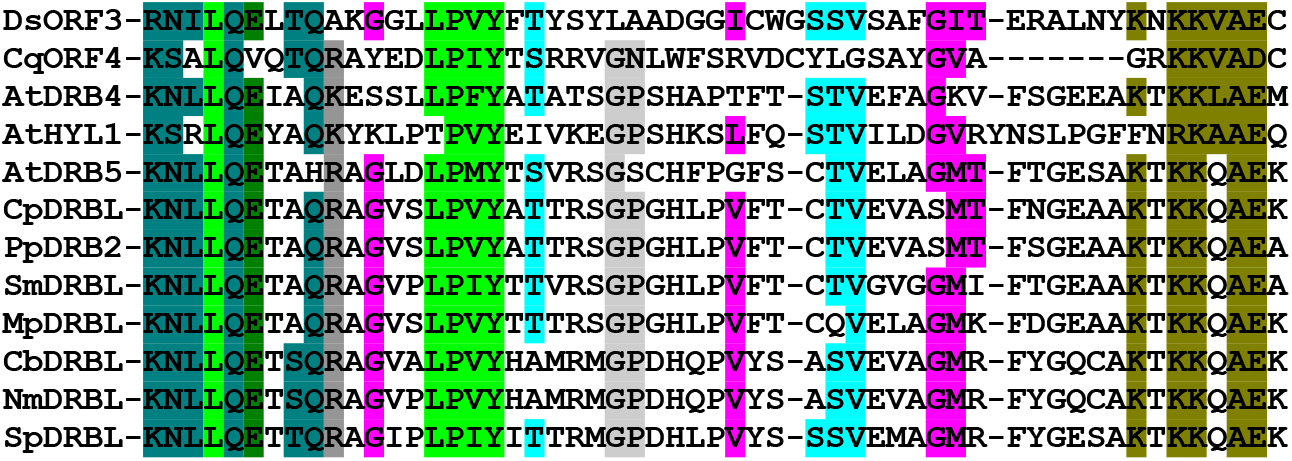
Multiple sequence alignment of the dsRNA-binding centers of proteins HYL1, DRB4 and DRB5 from *A. thaliana* with vDRB proteins of *D. scoparium* (DsORF3) and *C. quitensis* (CqORF4) as well as DRB-like proteins of moss *Ceratodon purpureus* (CpDRBL - accession KAG0625911), moss *Physcomitrium patens* (PpDRB2 - XP_024393530), lycophyte *Selaginella moellendorffii* (SmDRBL - EFJ14280), liverwort *Marchantia polymorpha* (MpDRBL - PTQ26790), charophyte algae *Chara braunii* (CbDRBL - GGXX01036972), charophyte algae *Nitella mirabilis* (NmDRBL - JV799478), charophyte algae *Spirogyra pratensis* (SpDRBL - GFWN01008525).

**Fig. 5.**
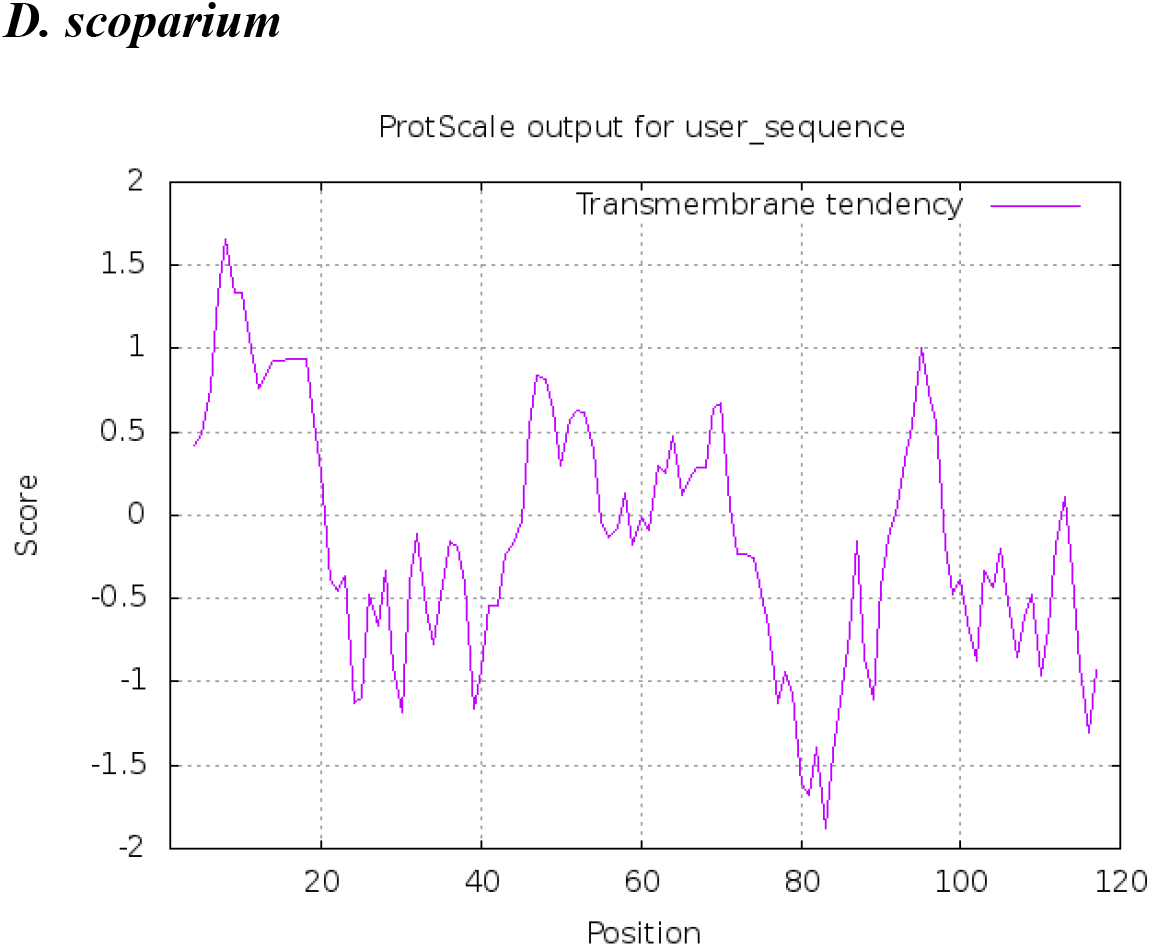
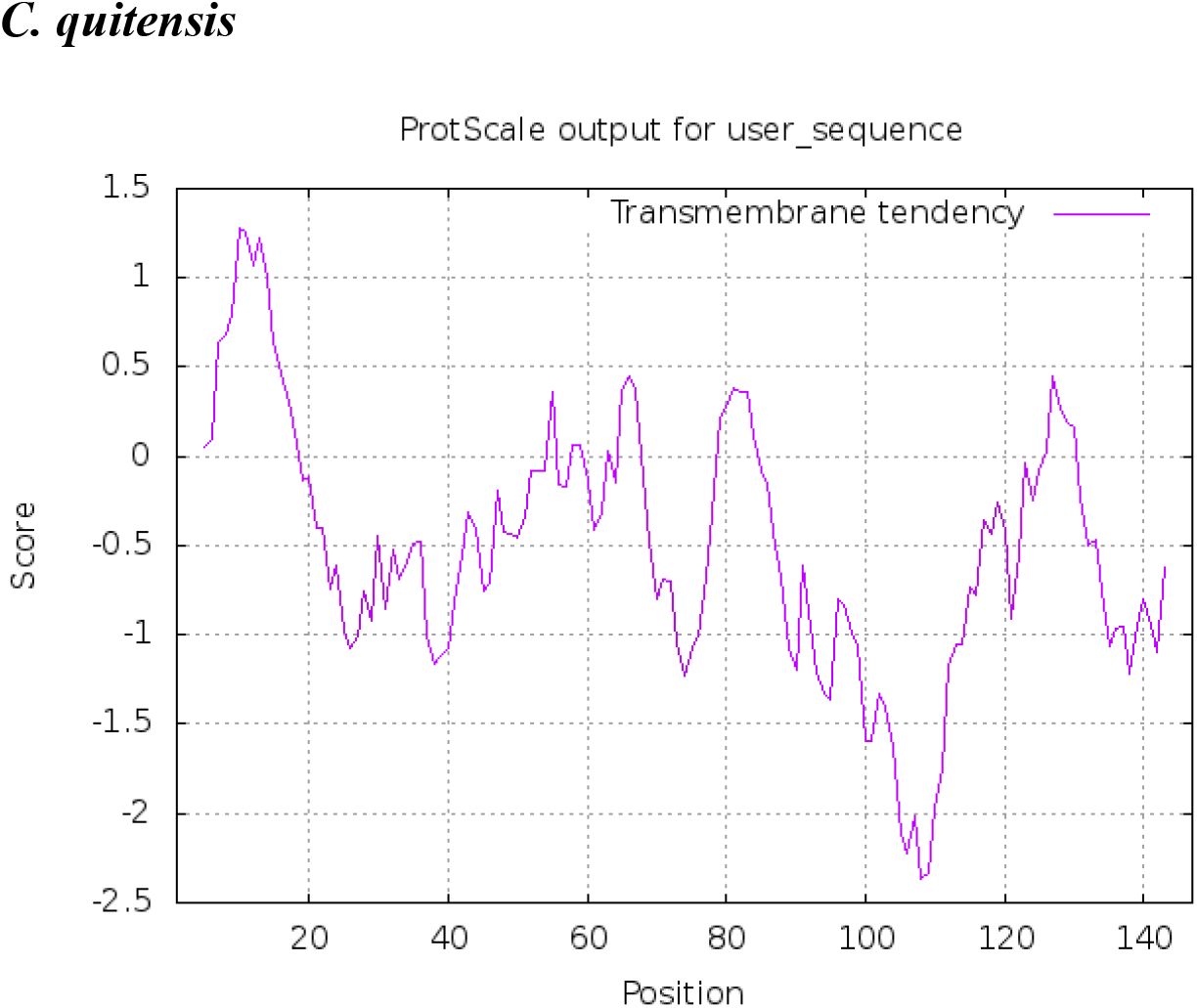
Prediction of hydrophobic membrane-bound regions (https://web.expasy.org/protscale/) in vDRB proteins of *D. scoparium* (DsORF3) and *C. quitensis* (CqORF4). Membrane-bound protein segments are above 0.8 cutoff line.

The ORF4 in Ds-VLRA2 overlaps the ORF3 by 60 nucleotides (Fig. 2A) and encodes protein of 361 residues containing motifs characteristic for helicases of SF1 superfamily and showing obvious similarity to BMB1 helicases (Fig. 1). ORF4 is followed by overlapping ORF5 and ORF6 (Fig. 2A), which code for small hydrophobic proteins with two putative transmembrane domains and central hydrophilic region related to TGB2/BMB2 proteins (Fig. 6). Protein sequence similarity of the encoded proteins and general organization of the gene block represented by ORFs 4-6 of Ds-VLRA2 resembles TGB and related Cq-TGB-like module (accession GCIB01126289).

**Fig. 6.**
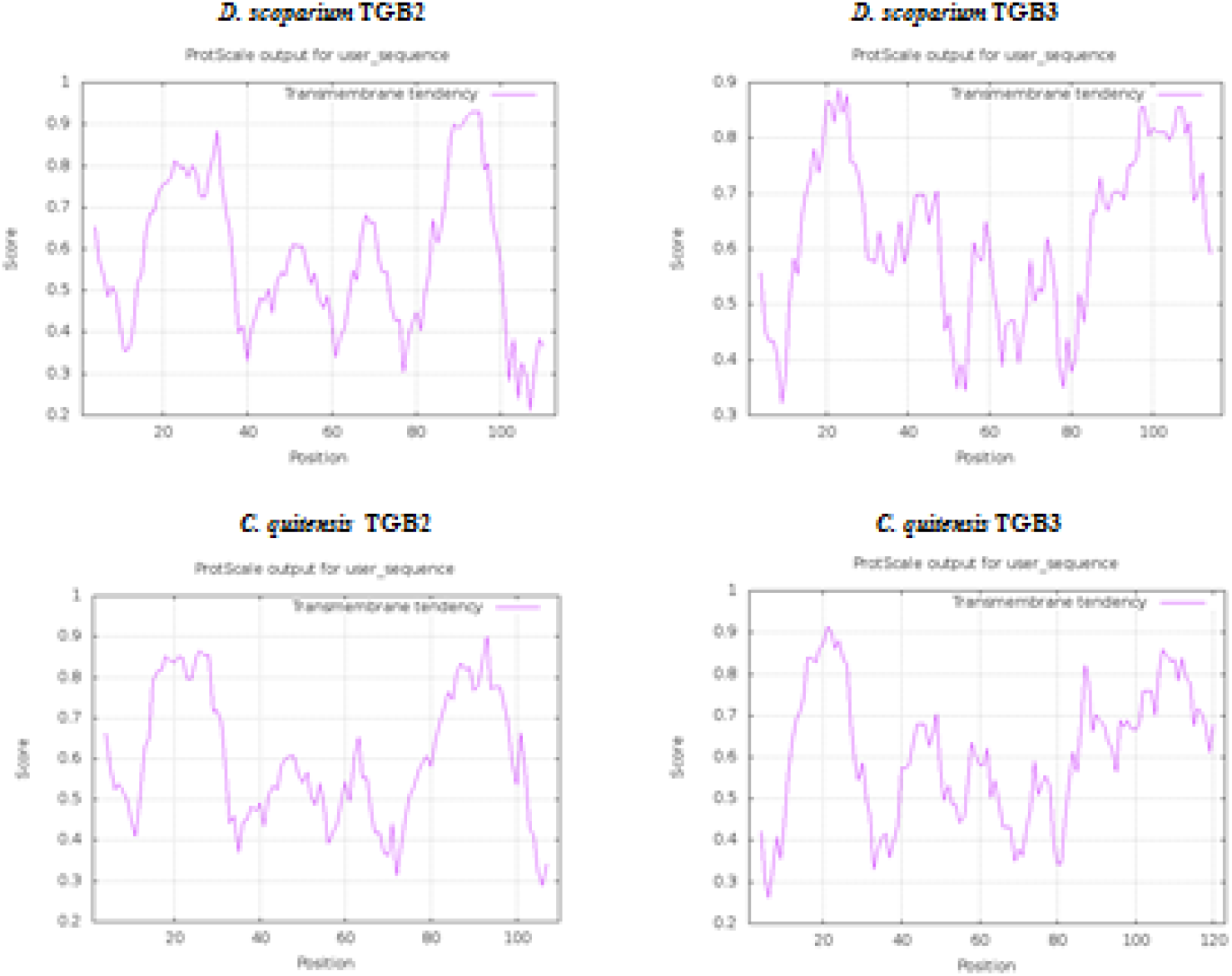
Prediction of hydrophobic membrane-bound regions (https://web.expasy.org/protscale/) in TGB-like proteins of *D. scoparium* and *C. quitensis* VLRAs.

Taking into account the similarity of Ds-VLRA2 TGB and Cq-TGB, we further analyzed whether the previously assembled *C. quitensis* TGB-containing VLRA was incomplete and could be extended into the 5’-terminus direction. Our *de novo* assembly of transcriptome sequencing data for *C. quitensis* SRA experiment (SRX814890) resulted in the probable full-length contig (Cq-VLRA2) of 4,718 nt in length excluding the poly(A) tail. The resulting sequence was confirmed using incomplete contigs from TSA database (GCIB01147888, GCIB01142942, GCIB01133924 and GCIB01126289). The ORF prediction at ExPASy showed that this contig contains seven ORFs flanked by a 5’-untranslated region (5’-UTR, at least 259 nt) and a 3’-UTR (213 nt). The 5’-terminal ORF1 encodes a capsid protein (164 aa in length) (Fig 2A) showing similarity to the wheat stripe mosaic virus CP (genus *Benyvirus*) (YP009553316, 27% identity, E-value 1e-04), and the sorghum chlorotic spot virus CP (genus *Furovirus*; family *Virgaviridae*) (NP659022, 29% identity, E-value 3e-04). The next ORF2 represents read-through domain of CP fusion protein as it was reported for benyviruses (Fig. 2B) (Saito et al., 1996). Among three described types of read-through nucleotide signatures, ORFs1/2 contain the type I motif containing a UAG codon, which is followed by the consensus motif CARYYA (where R is a purine and Y is a pyrimidine). This mechanism of translation is also used in tobamovirus replicase genes (Firth and Brierley, 2012; Miras et al., 2017).

ORF2 is followed by an intergenic region of 73 nucleotides in length and an ORF3, which codes for a small protein with the charged N-terminal half and cysteine-rich C-terminal region (Fig. 2A, Fig. 7). This cysteine-rich protein (CRP) shows no sequence similarity to the RNA2-encoded benyvirus CRP (Fig. 2B), and its cysteine-rich region is marginally similar to double zinc finger motif-containing module of FYVE domain involved in mRNA transport to endosomes (Pohlmann et al., 2015).

**Fig. 7.**
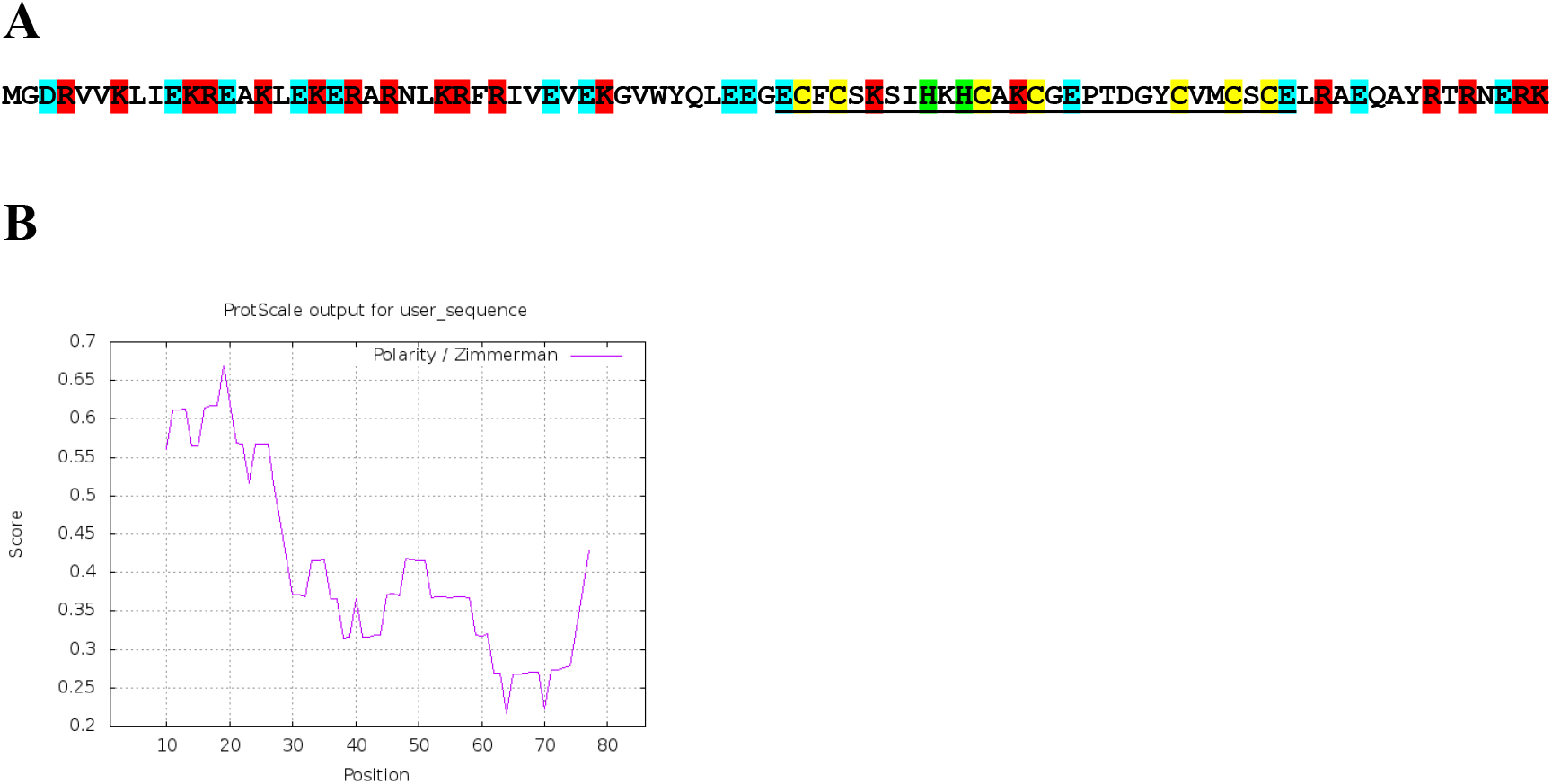
(**A**) Amino acid sequence of an ORF3 protein encoded by Cq-VLRA2. Positively charged residues are shown in red, negatively charged – in blue, cysteines – in yellow. Putative zinc finger motif-containing module is underlined. (**B**) Polarity plot of ORF3 protein (https://web.expasy.org/protscale/).

The NCBI BLAST analysis showed that Cq-VLRA2 ORF4 protein is quite similar to Ds-ORF3 (vDRB) protein and contains single DSRM with signatures specific for DSRM_AtDRB-like proteins (Fig. 4) and the N-terminal hydrophobic segment (Fig. 5). Cq-ORF4 is followed by an overlapping ORF5 encoding protein of 355 amino acids in length (Fig. 2A). The Cq-ORF5 protein (earlier named Cq-TGB1, see above) possesses motifs characteristic for helicases of SF1 superfamily and shows significant similarity to Ds-ORF4 protein (Ds-TGB1) and BMB1 helicases (Fig. 1). Like Ds-VLRA2, Cq-ORF4 is followed by overlapping ORFs 5 and 6 (Fig. 2) encoding small hydrophobic proteins with two putative transmembrane domains and central hydrophilic region related to Ds-ORF5/6 proteins (Fig. 6).

A general view on the organization of Ds-VLRA2 and Cq-VLRA2 strongly suggests two conclusions: 1) Both virus-like RNAs have considerable similarity to the benyvirus TGB-containing RNA2. Indeed, RNA2 of the type benyvirus Beet necrotic yellow vein virus (BNYVV) has six ORFs, namely, the CP gene terminated by a leaky stop codon, the CP read-through protein gene, the TGB and the cistron coding for a cysteine-rich protein having a silencing suppressor activity (Fig. 2B) (Saito et al., 1996; Chiba et al., 2013); 2) These RNAs include a conserved module of four overlapping genes, which is proposed to be named “Tetra-Cistron Movement Block” (TCMB). In comparison with the TGB and BMB modules, TCMB includes an additional 5’-terminal ORF, which overlaps the downstream gene and codes for the vDRB protein with a novel, previously undescribed for virus-encoded proteins, dsRNA-binding activity. Importantly, the cellular DRBs were shown to be incorporated into virus-specific replication membrane compartments (Barton et al., 2017; Incarbone et al., 2021), the structures often located at the PD orifice and involved in virus cell-to-cell movement (Tilsner et al., 2013). Similarly, the related hydrophobic vDRB proteins could be proposed to work in concert with other TCMB proteins to take part in viral dsRNA delivery to and/or retaining in PD-associated ER membrane-derived replicative compartments (Tilsner et al., 2013; Lazareva et al., 2021) and, thus, participate in virus cell-to-cell movement.

### PROPOSED GENERAL ORGANIZATION OF TCMB-CONTAINING PLANT VIRUS GENOMES

Assuming similarity of Ds-VLRA2 and Cq-VLRA2 to benyvirus RNA2 (Fig. 2) in gene organization, we performed search of the NCBI *Dicranum scoparium* TSA database in an attempt to find Ds-RNA1 expecting to code for virus replicase as in the case of BNYVV. As an initial query, we used 150 amino acid-long segment of BNYVV replicase (GDD domain). BLAST search revealed a single TSA contig (HANF01090670) of 305 nucleotdes in length which codes for a protein segment containing a GDD motif typical for RNA-dependent RNA polymerase (RdRp) domain and having more than 60% protein identity to BNYVV replicase protein (data not shown). To assemble the expected Ds-RNA1, transcriptome sequencing data for *D. scoparium* SRA experiment ERX3824048 linked to the TSA project were used. The assembled full-length sequence of the contig comprised 6,624 nt, excluding the poly(A) tail. ORF prediction showed that the contig contains a single cistron encoding viral replicase flanked by a 5’-UTR (at least 78 nt) and a 3’ UTR (237 nt) (Fig. 8A). Importantly, pairwise BLASTN analysis of the 3′-untranslated regions from Ds-VLRA1 and Ds-VLRA2 indicated a significant degree of sequence conservation among them and strongly suggested that both moss VLRAs are indeed the two components of a single virus genome (Fig. 8B).

**Fig. 8.**
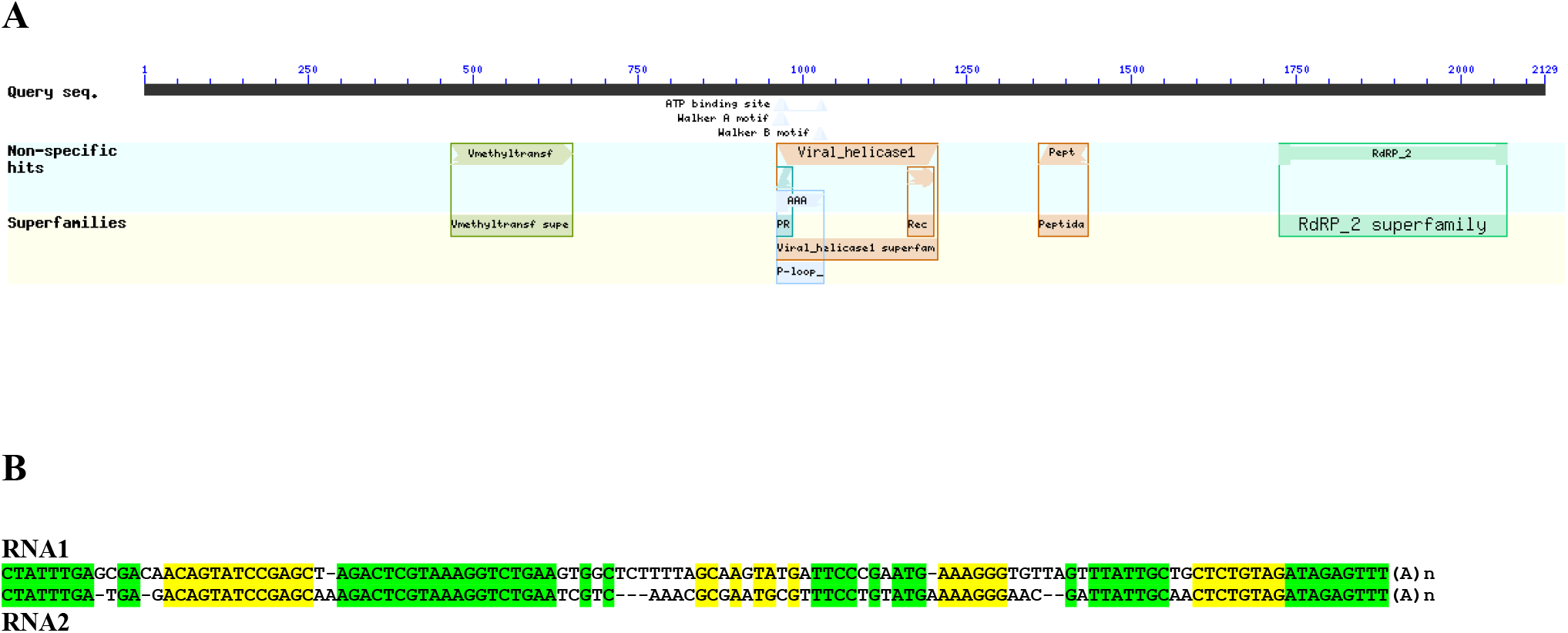
(**A**) The predicted organization of ORF1 protein of Ds-VLRA1 (NCBI formate) containing four conserved domains: a viral methyltransferase domain (Vmethyltransf, amino acid positions 440–625); a viral helicase 1 domain (amino acid positions 939-1179); papain-like proteinase domain (Pept, positions 1333-1408), and RdRp core motif (RdRP 2 superfamily, amino acid positions 1698–2045). (**B**) Nucleotide sequence alignment of the 3′-terminal regions preceding poly(A) in the predicted VLRA RNAs 1 and 2 from *Dicranum scoparium*. Highly conserved RNA blocks are highlighted by yellow and green background.

ORF1 protein of Ds-VLRA1 contains four conserved domains: a viral methyltransferase domain (MTR, pfam01660, amino acid positions 440–625, E-value 2.58e-06); a viral helicase 1 domain (HEL, pfam01443, amino acid positions 939-1179, E-value 4.70-22); papain-like proteinase domain (PROT, pfam05415, positions 1333-1408, E-value 6.97-06), and RdRp core motif (pfam00978, amino acid positions 1698–2045, E-value 2.42e-14) (Fig. 8A). The MTR domain is known to be conserved in a wide range of single-stranded RNA viruses, including orders *Martellivirales, Tymovirales and Hepelivirales* (Rozanov et al., 1992). All replicases in the members of these orders also encode HEL and RdRp domains containing typical motifs (Koonin and Dolja, 1993) conserved also in the ORF1 protein of Ds-VLRA1. Although the protease domain is not common for the above-mentioned replicases, Ds-VLRA1 encodes a protein domain with similarity to benyvirus protease (Fig. 8A), which is conserved in most benyviruses and required to produce mature replicase proteins by proteolytic self-cleavage (Rodamilans et al., 2018).

We further used encoded amino acid and nucleotide sequences of Ds-VLRA1 as queries for searches of *C. quitensis* SRA data (SRX814890) in order to assemble a Cq-RNA1 complete nucleotide sequence. However, only a rather short nucleotide sequence encoding a part of RdRp domain (including the GDD signature), which showed significant similarity to Ds-VLRA1 protein and moderate similarity to benyvirus replicases, has been assembled (Fig. 9).

**Fig. 9.**
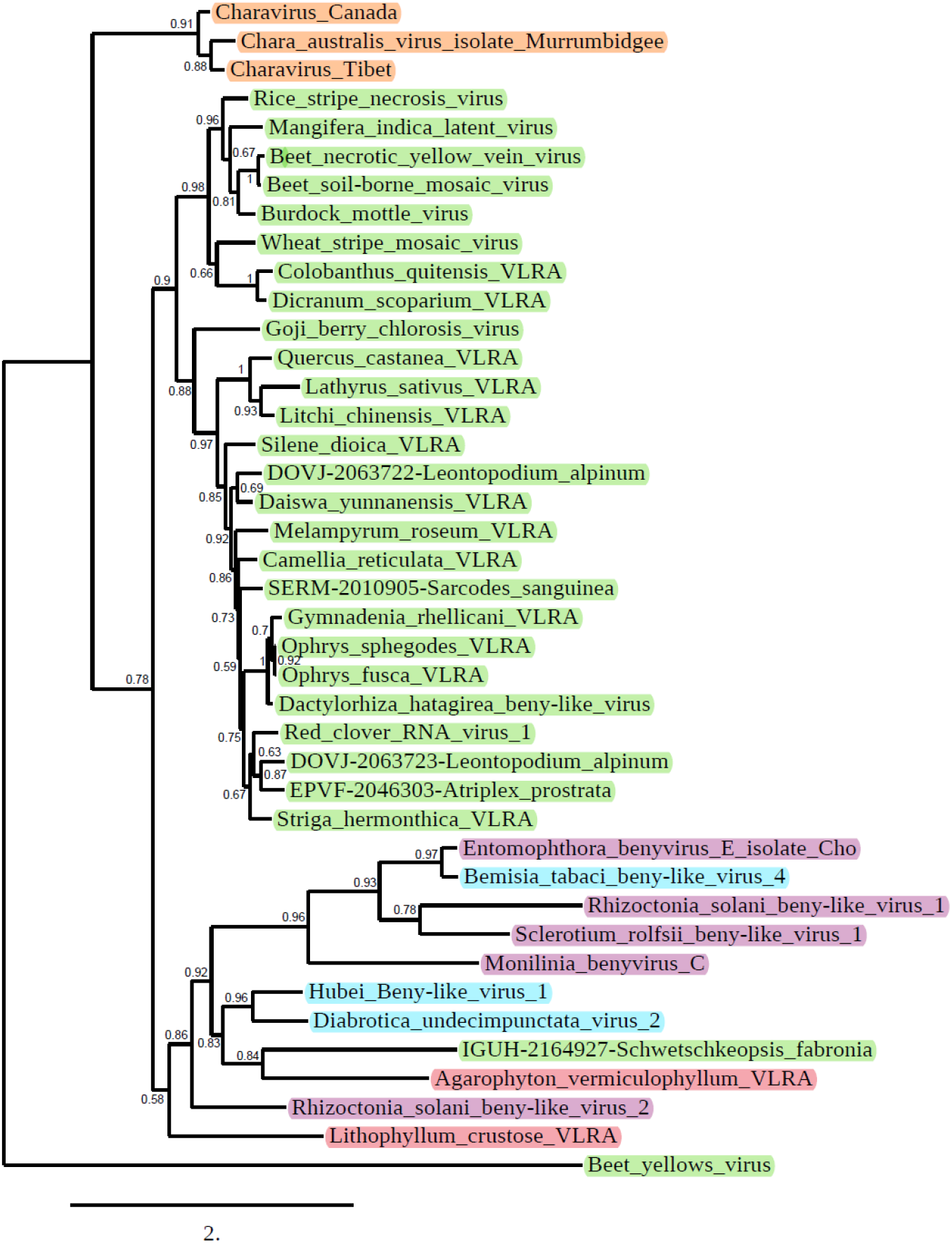
Phylogenetic analysis of the conserved motifs of RdRp derived from the aligned deduced amino acid sequences of beny-like viruses and selected VLRAs. The phylogenetic tree was constructed using the maximum likelihood method at Phylogeny.fr. The beet yellows closterovirus RdRp was used as outgroup. The bootstrap values obtained with 1000 replicates are indicated on the branches, and branch lengths correspond to the branch line’s genetic distances. The genetic distance is shown by the scale bar at the lower left. Charaviruses are shown in brown, plant viruses are in green, fungal viruses are in pink, arthropod viruses are in blue, red algae viruses are in rose.

### CLUES TO THE POSSIBLE ORIGINATION OF HELICASES ENCODED BY BMB AND TCMB

An important clue to the pathways of evolutionary origin of TGB, BMB and TCMB, to our mind, is provided by phylogenetic analysis of their encoded helicases. Evidently, BMB and TCMB helicases form a common branch which is closer to benyvirus TGB helicases and less similar to potex- and hordei-like TGB helicases (Fig. 1). Moreover, BMB and TCMB helicases show more sequence identity to beny-like replication helicases than to potex- and hordei-like TGB helicases (Morozov and Solovyev, 2015). Therefore, it can be suggested that a starting event in the evolutionary emergence of BMB- and TCMB could be duplication and autonomization of the replicative helicase domain occurred due to template switching during the virus genome replication along with probable non-replicative joining of RNA fragments (Bujarsky, 2013). Such RNA-RNA recombination likely resulted in the formation of the earliest monopartite and/or multipartite beny-like replicons with an autonomized copy of SF1 helicase. Recombination-dependent scenarios for evolutionary radiation have been also proposed for viruses of the family *Hepeviridae* (Kelly et al., 2016), which, together with benyviruses and tetraviruses (Dorrington et al., 2020), comprise the order *Hepelivirales* (Koonin et al., 2020). Taking into account the fact that the currently revealed TCMB-containing viruses infect primitive land plant (moss) (this paper) or the geographically long-term isolated flowering plant *C. quitensis* (Solovyev and Morozov, 2017), TCMB might be considered as an evolutionary old movement gene module originated in benyvirus-like replicons.

### ORIGINATION OF HYDROPHOBIC PROTEIN GENES IN MOVEMENT GENETIC MODULES

As it was suggested above, possible starting event in evolution of BMB- and TCMB-containing viruses was the formation of the beny-like replicons with an autonomized (duplicated) copy of beny-like SF1 replicative helicase. However, phenomenon of origination and acquisition of hydrophobic protein genes in different types of movement gene modules is generally obscure (Morozov and Solovyev, 2015; 2020). In this study, we found that among monopartite plant beny-like viruses, in addition to BMB-containing replicons, many VLRAs and viruses contain one or two small ORFs, which are located downstream of the replicase gene and encode small “orphan” proteins with one or two hydrophobic segments (Fig. 9 and Fig. 10) (Solovyev and Morozov, 2017).

**Fig. 10.**
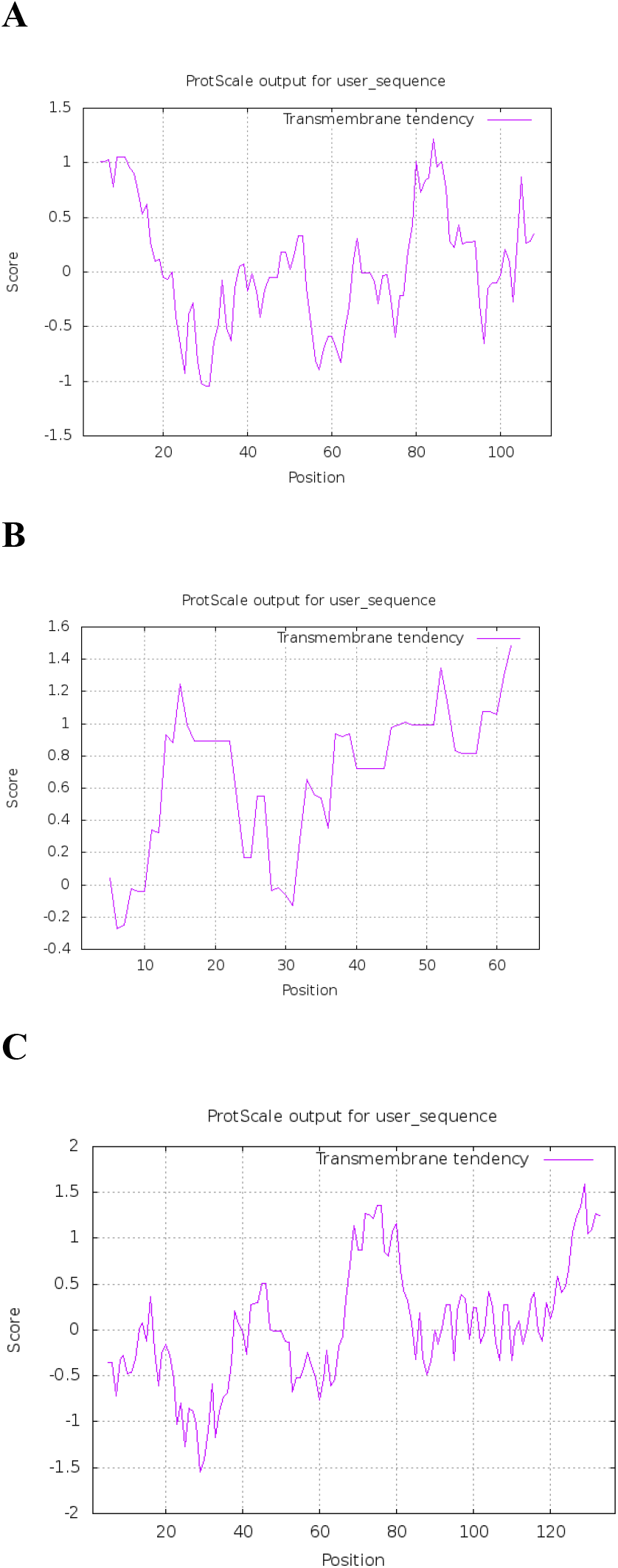
Prediction of hydrophobic membrane-bound regions (https://web.expasy.org/protscale/) in p2 (**A**) and p3 (**B**) “orphan” proteins of Red clover RNA virus 1 (accession MG596242) as well as ORF2 protein (**C**) of Dactylorhiza hatagirea beny-like virus (accession BK013327). Membrane-bound protein segments are above 0.8 cutoff line.

We hypothesize that these hydrophobic “orphan” protein ORFs have originated due to recombination of viral RNAs with host transcripts containing *de novo* emerged ORFs. Indeed, it was proposed that novel eukaryotic “orphan” protein-coding genes can arise *de novo* in non-coding sequences, which, thus, may serve as a continuous reservoir of variable novel polypeptides serving as a raw material for natural selection (Vakirlis et al., 2020). Moreover, evolution of non-coding thymine-rich sequences can result in preferable emergence of ORFs encoding proteins with hydrophobic domains (Vakirlis et al., 2020; Fesenko et al., 2021). These findings allowed our colleagues to propose a novel evolutionary model suggesting that ORFs for small membrane-bound polypeptides emerging *de novo* in basal land plants could be a preferential subject of adaptive evolution because of escape of their encoded proteins from degradation or other deleterious interactions in the membrane environment (Fesenko et al., 2021). Therefore, viruses similar to the above-mentioned monopartite beny-like replicons, which might be produced through recombination with mRNAs carrying such ORFs for membrane polypeptides, can be proposed to serve as sources of ancestral membrane protein genes for recombination-dependent evolution TCMB- and BMB-like genetic modules and also some other movement genetic modules described for plant viruses (Morozov and Solovyev, 2020).

### EVOLUTIONARY RADIATION OF BENYVIRUS-LIKE RNA REPLICONS

Assuming the proposed involvement of replicative beny-like helicase in evolution of the above-mentioned movement genetic modules, it is tempting to speculate on the global evolution of beny-like replication proteins. Generally, a phylogenetic tree of selected beny-like RdRp domains (Fig. 9) showed three main branches including (i) a basal branch composed of closely related RNA viruses from fresh-water species of algal genus *Chara found in Australia and Canada* (Gibbs et al., 2011; Vlok et al., 2019); (ii) a branch of benyviruses (Niehl et al., 2020), plant bipartite viruses with TCMB (this study) and related monopartite plant beny-like viruses and VLRAs; (iii) a mixed branch including fungal and arthropod viruses, as well as VLRAs from red algae. Indeed, recent advances in sequencing benyvirus-like RNA replicons revealed their multiple hosts not only in plants but also among arthropods (particularly, Bemisia tabaci beny-like virus 6 - MW256699; Bemisia tabaci beny-like virus 4 - MW256697; Hubei Beny-like virus 1 - MK231108; Diabrotica undecimpunctata virus 2 - MN646771) and fungi (particularly, Erysiphe necator associated beny-like virus 1 - MN617775; Rhizoctonia solani beny-like virus 1 - MK507778; Sclerotium rolfsii beny-like virus 1 - MH766487) (Fig. 9) (Shi et al., 2016; Zhu et al., 2018; Picarelli et al., 2019; Gilbert et al., 2019; Liu et al., 2020).

Significantly, among monopartite Viridiplantae viruses, the beny-like replicase is encoded by two viruses with an unusual gene organization, namely, beny-like Chara virus (Vlok et al., 2019) and goji berry chlorosis virus (GBCV) (Kwon et al., 2018). It is known that charophyte algae are considered as the ancestors of land plants, and Chara viruses may be evolutionarily related to ancestor virus species that infected first plants colonizing terrestrial habitats (Vlok et al., 2019). Interestingly, beny-like Chara viruses are distributed around the globe, since in addition to species found in Australia and North America we revealed closely related viral RNA metagenomic sequences in the NCBI Sequence Read Archive (SRX8007769), BioProject accession PRJNA615325 (data not shown). These data were derived from samples of fish gills collected from Qinghai Lake in Tibet. The largest encoded protein of these monopartite viruses shows the relationship with RNA polymerases of benyviruses (Fig. 9), while the capsid protein is distantly related to the tobamovirus CP (Gibbs et al., 2011; Vlok et al., 2019). Two additional open reading frames (ORFs) code for an RNA helicase and a protein of unknown function. Importantly, this non-replicative “accessory” helicase is related to helicases of SF-2 superfamily in contrast to “accessory” TGB1 helicases belonging to SF-1 (Vlok et al., 2019). It is clear the replicase and tobamo-like CP genes belong to different lineages of the alphaviruses, orders *Hepelivirales* and *Martellivirales*, respectively, whereas the “accessory” helicase to replicative helicase of viruses belonging to order *Amarillovirales* (Vlok et al., 2019).

The GBCV genome encodes six polypeptides. Strikingly, the replicase (ORF1) is more similar to benyvirus-like replicases, whereas coat protein (ORF2) is more closely related tobamovirus-like CPs (Solovyev and Makarov, 2016) and ORF5 encodes a movement protein related to the tobamovirus-like MP. Unusual genome organization suggests that GBCV may represent a recombinant between the viruses from families *Benyviridae* and *Virgaviridae* (Kwon et al., 2018). This evolutionary episode also suggests a realistic possible pathway for advanced evolution of TGB, where the horizontal gene transfer of this gene module from beny-like RNA replicons could occur to ancestral replicons belonging to viruses belonging to orders *Martellivirales and Tymovirales* and initiate the evolution of potex-like and hordei-like TGBs.

Recent studies suggest not only ways of radiation of genome organization for beny-like replicons but also approximate gene divergence dates. It was shown that the estimates of gene divergence dates for the RdRp and CP proteins from *Virgaviridae* and *Benyviridae* are quite different. Generally, wide distribution of tobamo-like CP genes (Solovyev and Makarov, 2016) in viruses of orders *Hepelivirales* and *Martellivirales* strongly suggests significant role of horizontal gene transfer in evolutionary radiation of these genes (Shi et al., 2016). The divergence of the charavirus CP with that of tobamoviruses (family *Virgaviridae*) was estimated to be 212 million years ago (mya) (Vlok et al., 2019). On the other hand, time for divergence between charavirus/benyvirus RdRp (order *Hepelivirales*) and virga-like RdRp genes (order *Martellivirales*) was estimated to be ∼900 mya (Vlok et al., 2019). So, it seems that benyvirus-like replicases started their evolutionary radiations in late Precambrian, *i*.*e*. perhaps even before the chlorophyte-charophyte split likely occurred 850–1100 million years ago (Del Cortona et al., 2020; Strassert et al., 2021). In this respect, it is important that the red algae (Rhodophyta) are most ancient in the kingdom Plantae (Archaeplastida) (https://www.algaebase.org/browse/taxonomy/), and an origin of multicellular red algae is expected around 1000-1600 mya (Schön et al., 2021; Carlisle et al., 2021; Strassert et al., 2021). So, the divergence between replication proteins of viruses in orders *Hepelivirales* and *Martellivirales* could occur in marine red algae species. In support for the proposed role of Rhodophyta as a host for common ancestors of *Hepelivirales* and *Martellivirales*, it was found that the both types of virus replicons can be found in modern red algae hosts (Fig. 11).

**Fig. 11.**
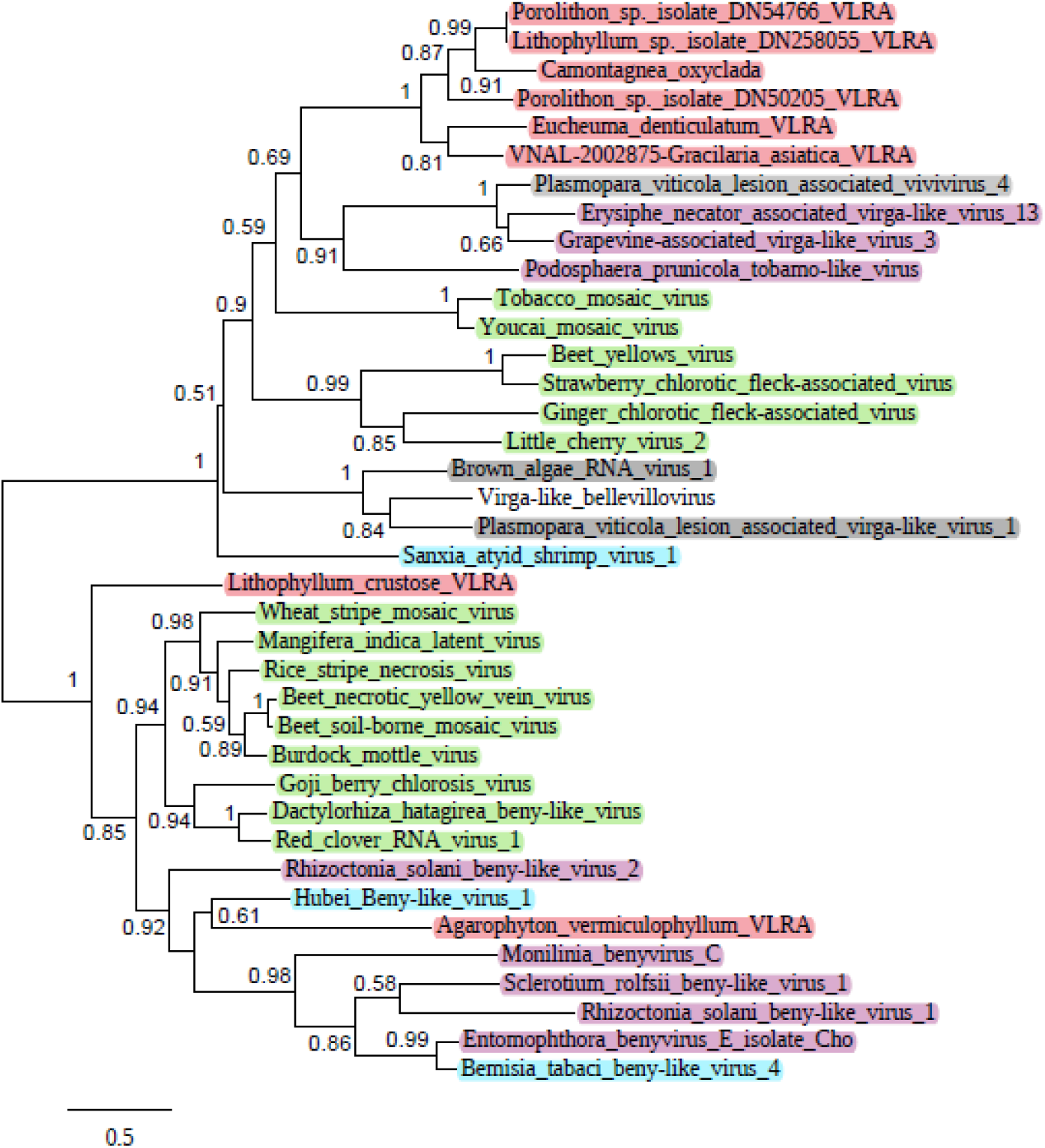
Phylogenetic analysis of the conserved motifs of RdRp derived from the aligned deduced amino acid sequences of selected beny-like replicative proteins (bottom branch), virga-like replicative proteins (upper branch) and red algae VLRAs. The unrooted phylogenetic tree was constructed using the maximum likelihood method at Phylogeny.fr. The bootstrap values obtained with 1000 replicates are indicated on the branches, and branch lengths correspond to the branch line’s genetic distances. The genetic distance is shown by the scale bar at the lower left. Viruses of Stramenopiles are shown as shaded, plant viruses are in green, fungal viruses are in pink, arthropod viruses are in blue, red algae viruses are in rose.

### EXPERIMENTAL

Virus nucleotide and protein sequences were collected from the NCBI database. Assembled viral genomes were mainly extracted from NCBI database. The sequence comparisons were carried out using the BLAST algorithm (BLASTn and BLASTp) at the National Center for Biotechnology Information (NCBI). Open reading frames (ORFs) were identified using the NCBI ORF Finder program (http://www.bioinformatics.org/sms2/orf_find.html). Gene translation and prediction of deduced proteins were performed using ExPASy (http://web.expasy.org/translate/). Conserved motif searches were conducted CDD (http://www.ncbi.nlm.nih.gov/Structure/cdd/wrpsb.cgi) databases. Coiled-coil protein regions were predicted using Waggawagga software (https://waggawagga.motorprotein.de/) (Simm et al., 2021).

To assemble the full-length plant VLRAs, transcriptome sequencing data for *D. scoparium* and *C. quitensis* SRA experiments linked to the TSA projects were downloaded using fastq-dump tool of NCBI SRA Toolkit 2.9.0. (http://ncbi.github.io/sra-tools/). Reads quality was checked with FastQC (https://www.bioinformatics.babraham.ac.uk/projects/fastqc/). *De novo* assembly of VLRAs coding for TCMB modules was carried out using SPAdes 3.12.0 (Bankevich et al., 2012) in “RNA mode”.

Phylogenetic analysis was performed with “Phylogeny.fr” (a free, simple to use web service dedicated to reconstructing and analysis of phylogenetic relationships between molecular sequences) by constructing maximum likelihood phylogenetic trees (http://www.phylogeny.fr/simple_phylogeny.cgi). Bootstrap percentages received from 1,000 replications were used.

## ACKNOWLEDGMENTS

The authors are grateful for the funding received by the Russian Foundation for Basic Research (grant 20-04-00456).

## AUTHOR CONTRIBUTIONS

SM collected and analyzed the data, authored drafts of the paper;

AS authored drafts of the paper, prepared figures, reviewed the final draft.

## CONFLICT OF INTEREST STATEMENT

The authors declare that the research was conducted in the absence of any commercial or financial relationships that could be construed as a potential conflict of interest.

## DATA AVAILABILITY STATEMENT

The datasets presented in this study can be found in online repositories. The names of the repository/repositories and accession number(s) mentioned in this paper can be found at: https://www.ncbi.nlm.nih.gov/.

## Notes

https://www.ncbi.nlm.nih.gov/

### Competing Interest Statement

The authors have declared no competing interest.

### Summary of Updates

abstract updated author acknowledgment updated

